# Rigid DNA nanotube tethers suppress high frequency noise in dual-trap optical tweezers systems

**DOI:** 10.1101/2021.11.15.468716

**Authors:** Alan Shaw, Rohit Satija, Eduardo Antunez de Mayolo De la Matta, Susan Marqusee, Carlos Bustamante

## Abstract

Dual trap optical tweezers are a powerful tool to trap and characterize the biophysical properties of single biomolecules such as the folding pathways of proteins and nucleic acids, and the chemomechanical activity of molecular motors. Despite its vastly successful application, noise from drift and fluctuation of the optics, and Brownian motion of the trapped beads still hinder the technique’s ability to directly visualize folding of small biomolecules or the single nucleotide stepping of polymerases, especially at low forces (<10 pN) and sub-millisecond timescales. Rigid DNA nanotubes have been used to replace the conventional dsDNA linker to reduce optical tweezers noise in the low force range. However, optical tweezers are used to study a wide range of biophysical events, with timescales ranging from microseconds to seconds, and length changes ranging from sub nanometers to tens of nanometers. In this study, we systematically evaluate how noise is distributed across different frequencies in dual trap optical tweezers systems–and show that rigid DNA nanotube tethers suppress only high frequency noise (kHz), while the low frequency noise remains the same when compared to that of dsDNA tethers.

## Introduction

Recent advances in optical tweezers instrumentation have settled on a “dual trap” design to reduce mechanical drift in the system^1,2^. This setup uses two separate laser beams to trap one micron sized bead each, and the single biomolecule of interest is tethered in between the two beads with dsDNA as linker (Figure 1A, top panel). Both the Brownian motion of the trapped beads and the thermal fluctuation of dsDNA linkers introduce noise to the system and can obscure signals^2,3,4^. Noise reduction can be achieved by applying high force (>10 pN) to stiffen the dsDNA linker^5^, or by smoothing the raw data points^6^. Rigid DNA origami nanotubes have been applied to dual trap optical tweezers as rigid linkers and have been shown to reduce noise of the system, especially at low force (<10 pN)^3^. The authors showed that using their DNA nanotube linker system they can directly visualize the folding and unfolding transitions of a 6 nt DNA hairpin, while dsDNA linkers of the same length cannot. However, optical tweezers are used to study a wide spectrum of biophysical events, with timescales ranging from sub milliseconds to seconds, and distance changes ranging from sub nanometers to tens of nanometers^7–9^. To truly uncover the advantages of using rigid DNA nanotube tethers, we need to evaluate how optical tweezers noise is distributed across different frequencies, and how rigid DNA nanotube tethers alter this distribution. This will allow us to identify timescales where DNA nanotube tethers have maximum advantage in terms of noise suppression over dsDNA tethers.

**Figure 1:**
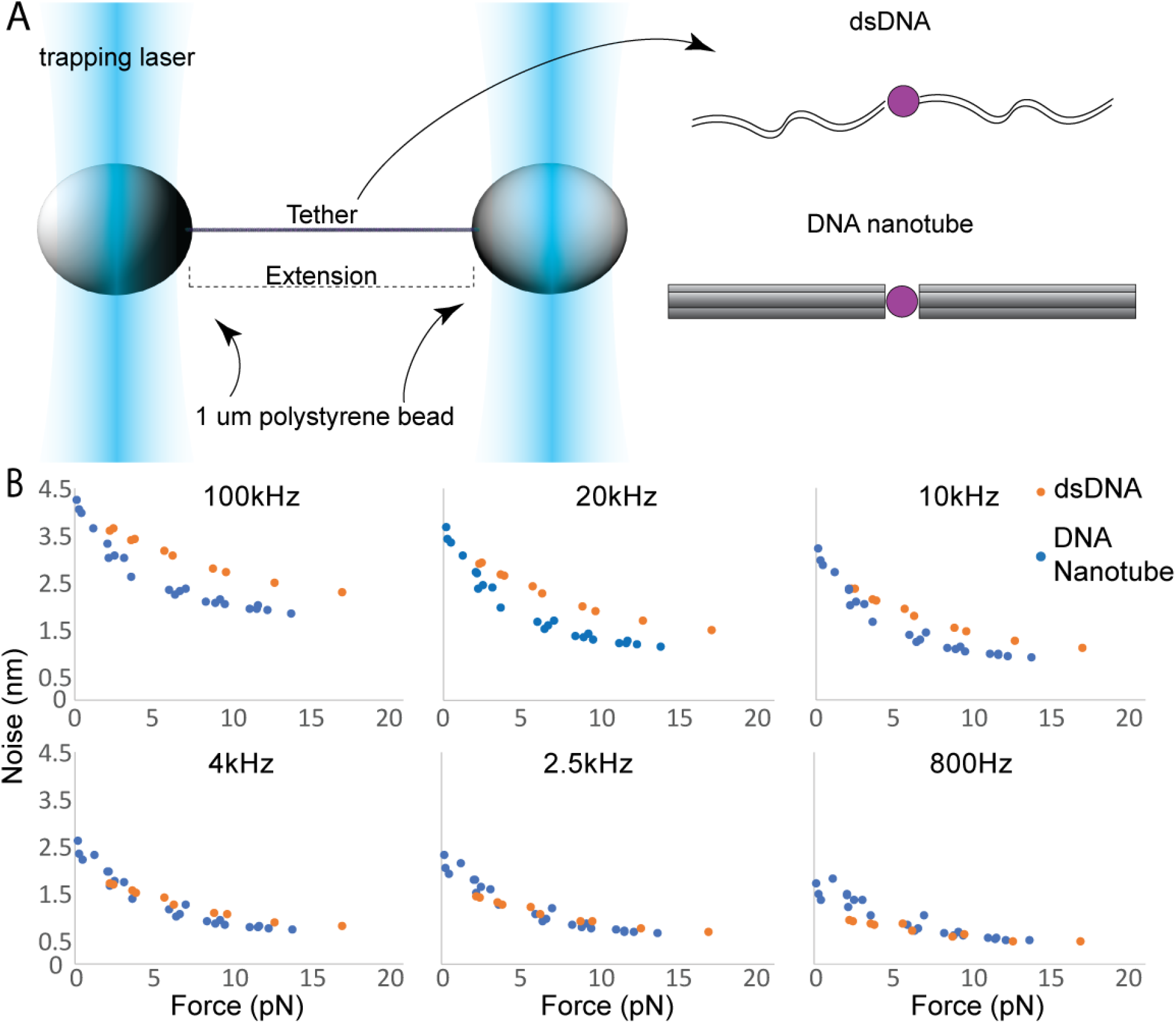
Noise suppression in dual trap optical tweezers using DNA nanotube tether. A) The dual trap optical tweezer setup uses two separate laser beams to trap two 1 um beads each, and a single tether between the beads was formed either using dsDNA or DNA nanotube. A single molecule of interest (purple circle), such as a DNA hairpin or protein, can be attached to the tether. The end-to-end distance of the tether (extension) is recorded. B) The noise in extension plotted against force applied by the optical tweezers at different sampling frequencies.

In this study, we designed and synthesized a 6-helix bundle DNA origami^10,11^ nanotube (Figure 1A, Supplemental Figure 1) with a design length of 1323 nt (449.8 nm). The choice of the 6-helix bundle was based on data presented by Pfitzner et al.^3^. Based on their data, the noise suppression effect of DNA nanotubes only became marginally better beyond the thickness of the 6-helix bundle. The nanotubes were assembled with either digoxigenin modified oligos or biotin modified oligos on one end and a complementary ssDNA overhang for dimerization on the other end (Supplemental figure 2). The digoxigenin and biotin modified nanotubes were first dimerized by DNA hybridization prior to loading onto anti-digoxigenin antibody modified 1 μm polystyrene beads. As a control, we used PCR with digoxigenin and biotin modified primers on lambda phage DNA to prepare dsDNA of the same length (2646 nt). Tethers were formed by bringing a streptavidin modified 1 um polystyrene bead close to the nanotube-loaded anti-digoxigenin bead, and the beads were quickly separated once a single tether was formed. Different forces were applied to a pair of beads with a single tether using passive mode^12^ (constant trap position), the extension between the two beads over time were recorded (Figure 1A), and the noise and its distribution across different frequencies of the measured extension were calculated (Figure 1B).

We discovered that DNA nanotube tethers only suppress high frequency noise, while low frequency noise remains at a similar level compared to that of dsDNA (Figure 1B). Moreover, analyzing the complete Power Spectral Density (PSD) of the experimental signal revealed anomalous frequency-dependence (Figure 2A), a hallmark of non-Markovian or memory effects in the dynamics. We modeled these dynamics using a generalized Langevin equation (GLE) framework, characterized by a friction memory kernel with a finite time decay, and showed, using exact analytical formulae and numerical simulations, how this memory can be exploited to reduce optical tweezers noise by varying the parameters of the beads, the traps, and the tether. Based on our analysis, we suggest that DNA nanotube tethers have an advantage over dsDNA tethers when studying fast transitions that occur in the sub-millisecond timescale, while there are no obvious advantages of using the nanotube tethers when studying events that occur in the millisecond timescale or slower. Using our DNA nanotube tethers, we show that we can directly visualize the fast-folding transition of a 6, 5, and 4 nt DNA hairpin. The folding of the 4 nt DNA hairpin is currently the smallest transition (about 2.5 nm) that occurs in the tens to hundreds of microseconds timescale that can be directly observed with optical tweezers.

**Figure 2:**
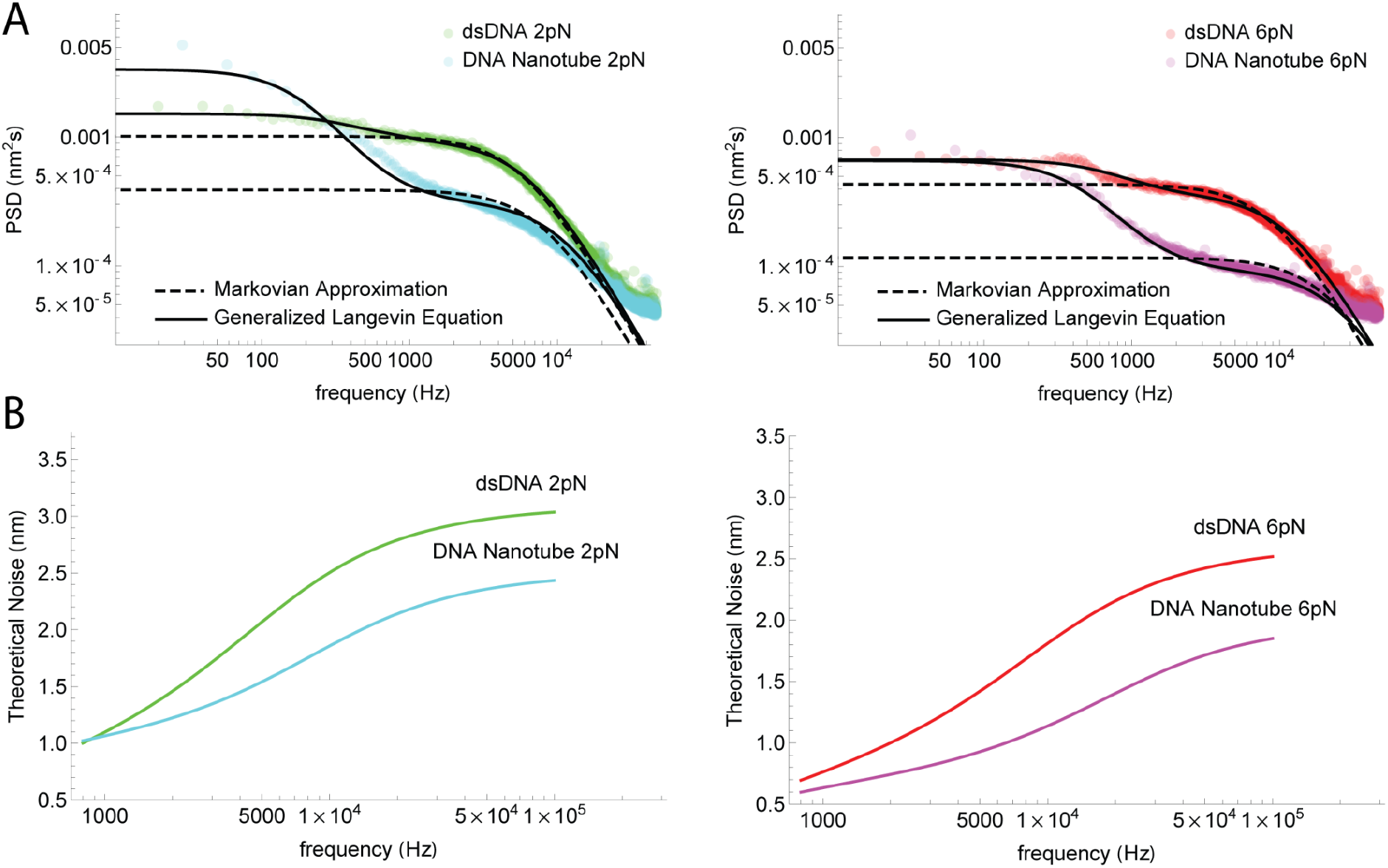
Comparison with theoretical models. A) Power Spectral Densities (PSD) for double stranded DNA and DNA Nanotube tethers held at different loads. The PSDs are either fit with the Generalized Langevin dynamics (Eq. 9) or its commonly used Markovian Approximation (Eq. 10). B) Theoretical noise,, 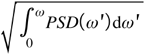 calculated using the GLE fits in the above plots plotted as a function of sampling bandwidth.

## Results

Successful assembly of the 6-helix bundle nanotube was confirmed by agarose gel electrophoresis and AFM imaging^13^ (Supplemental Figure 3, Supplemental Figure 4). The contour length of the nanotube was estimated using backbone tracing of the AFM images, which measured a contour length of about 447±15 nm. Force extension curves of the DNA nanotube tethers, when compared to that of dsDNA, show a sharper increase in force as it gets extended, and the force-extension curve of the control dsDNA fits well to the extensible worm-like chain model^14^ (Supplemental Figure 5). Both of these results corroborate well with previous studies^3,14^. Noise characterization was done by measuring the fluctuation in the extension between the two beads when the traps were held at constant positions. When we plot the noise (root-mean-squared extension) against the average force applied to the tethers, both tethers exhibit a monotonic decrease in noise as force increases (Figure 1B), which is the result of tensile stiffening of the tethers3. When comparing the noise difference between nanotube tethers and dsDNA tethers at 100 kHz sampling frequency, nanotubes clearly exhibit lower noise compared to dsDNA (Figure 1B top left panel), but the noise from the two tethers as the raw data is smoothed and downsampled to lower sampling frequencies became virtually identical below 4 kHz. This observation is peculiar, since a minimal model of 1D overdamped Langevin dynamics in a harmonic potential well predicts that increasing well curvature (trap stiffness) should decrease noise at all frequencies (Supplementary Figure 6).

On the contrary, projecting the dynamics of two linearly coupled microscopic beads onto the bead-to-bead extension indicates emergence of memory (see Appendix A for full derivation). Briefly, the standard model for dual optical trapping systems^15^ assumes that the dynamics are governed by the following set of coupled Langevin equations:

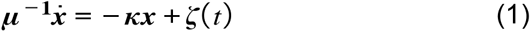

where 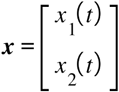 is a vector representing the positions of the two beads along the bead to bead coordinate, 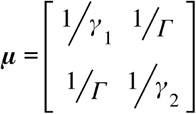 is the mobility matrix with *γ*_1_, *γ*_2_ representing the friction coefficients of the two beads and *Γ* representing the friction due to hydrodynamic coupling between them, 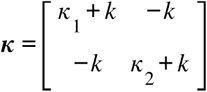 is the stiffness matrix corresponding to the linear conservative force, –*κx* = –∇*G*(*x*), due to the 2D potential:

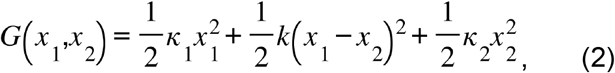

and the Gaussian distributed stochastic forces 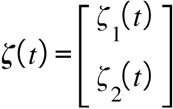 follow the correlation functions:

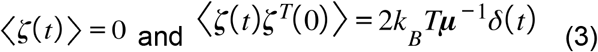

Here, *κ*_1_, *κ*_2_ are the stiffnesses of the two traps, *k*. is the stiffness of the tether (either dsDNA or DNA nanotube), *k_B_* is the Boltzmann constant, and *T* is the absolute temperature.

Since Eq. 1 is linear, we expect any linear combination of the bead positions, including the measured bead-to-bead extension *x*(*t*) = *x*_1_(*t*) – *x*_2_(*t*), to be described by a GLE^16^ of the form:

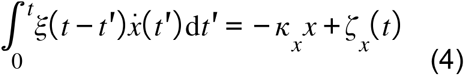

where *ξ*(*t*) is a time-dependent friction memory kernel, *κ_x_* is an effective spring constant along *x*, and *ζ_x_*(*t*) is a Gaussian-distributed random force with the following correlation functions:

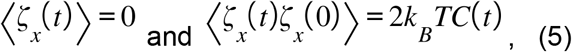

such that the memory kernel *ξ*(*t*) is related to *C*(*t*) in accordance with the fluctuation-dissipation theorem.

In Appendix A, we show that *ξ*(*t*) (or *C*(*t*)) in this model is of the form:

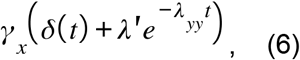

where the Markovian friction *γ_x_*, memory strength *λ′*, and memory time 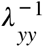 depend on the parameters *κ*_1_, *κ*_2_, *k*, *γ*_1_, *γ*_2_ and *Γ*. In particular, if the memory strength is too weak (*λ′* → 0), or the memory time is too short 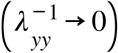, the non-Markovian effects become insignificant, i.e. *ξ*(*t*) = *C*(*t*) ≃ *γ_x_δ*(*t*), and Eq. 4 reduces to the overdamped Langevin equation in *x*:

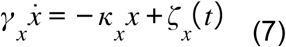

with the stochastic force having a white noise spectrum:

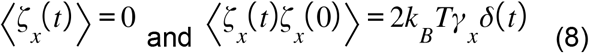

For the non-Markovian GLE in Eq. 4, the general form of the PSD in Fourier space can be derived as (Appendix B):

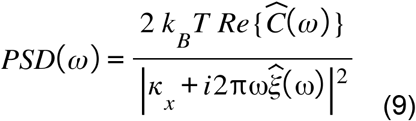

where for any function *f*(*t*), the Fourier Transform is defined as 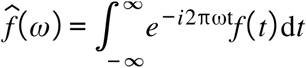 and the real part of the transform is 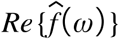. It is easy to see that in the Markov limit, Eq. 9 reduces to the well-known Lorentzian form:

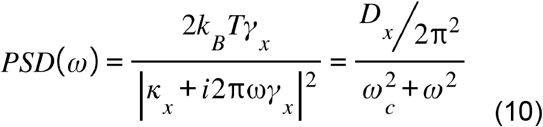

where 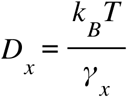 is the effective diffusion coefficient and 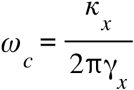 is the corner frequency along the bead to bead extension. In Figure 2A, we plot the PSD of our data using dsDNA and DNA nanotube tethers, along with fits to the GLE formula (Eq. 9) and its Markovian approximation (Eq. 10). We find that the Markovian approximation works well in case of dsDNA but fails to explain the anomalous frequency dependence observed in case of DNA nanotube tethers, whereas the GLE formula is able to explain both cases. Moreover, the noise at different bandwidths calculated using the GLE fits is consistent with the experimental observation that DNA nanotube tethers only suppress high-frequency noise (Figure 2B).

Based on our observation that DNA nanotube tethers only suppress high frequency noise, we suggest that the use of nanotube tethers have an advantage over dsDNA when studying biophysical events that occur in the sub-millisecond timescale. To demonstrate this, we introduced 6 nt, 5 nt, and 4 nt DNA hairpins to both nanotube and dsDNA tethers (Figure 3A) and the folding and unfolding of the hairpins were monitored under different loads. Existence of a single hairpin in the tether was confirmed by analyzing the force extension curves, which show a shoulder in the force range of 6 – 10 pN (Figure 3B left panel, dotted square). In addition, while noise decreases monotonically as tension increases when pulling on only the tethers, a bump in the monotonic decrease in noise as tension increases was observed when pulling on the tethers that contain hairpin (Figure 3B right panel, dotted square), suggesting that the tethered hairpin is frequently folding and unfolding within that force range. When analyzing data taken from passive mode and dsDNA tethers, Hidden Markov modeling^17^(HMM) failed to reliably capture the hairpin transitions between the folded and unfolded states, only yielding rates that are independent of the force applied (Figure 3C). When monitoring changes in hairpin extension using passive mode and nanotube tethers, two state hopping features of the hairpins can be clearly observed and the transitions can be captured with HMM (Figure 3D top panels). This allows us to extract the force-dependent folding and unfolding rates of the hairpins (Figure 3D bottom panels). The transition of the 4 nt hairpin is about 2.5 nm, and its folding and unfolding rates are in the range of 5 to 10 kHz. This is currently the fastest and smallest transition that is directly observable via dual trap optical tweezers.

**Figure 3:**
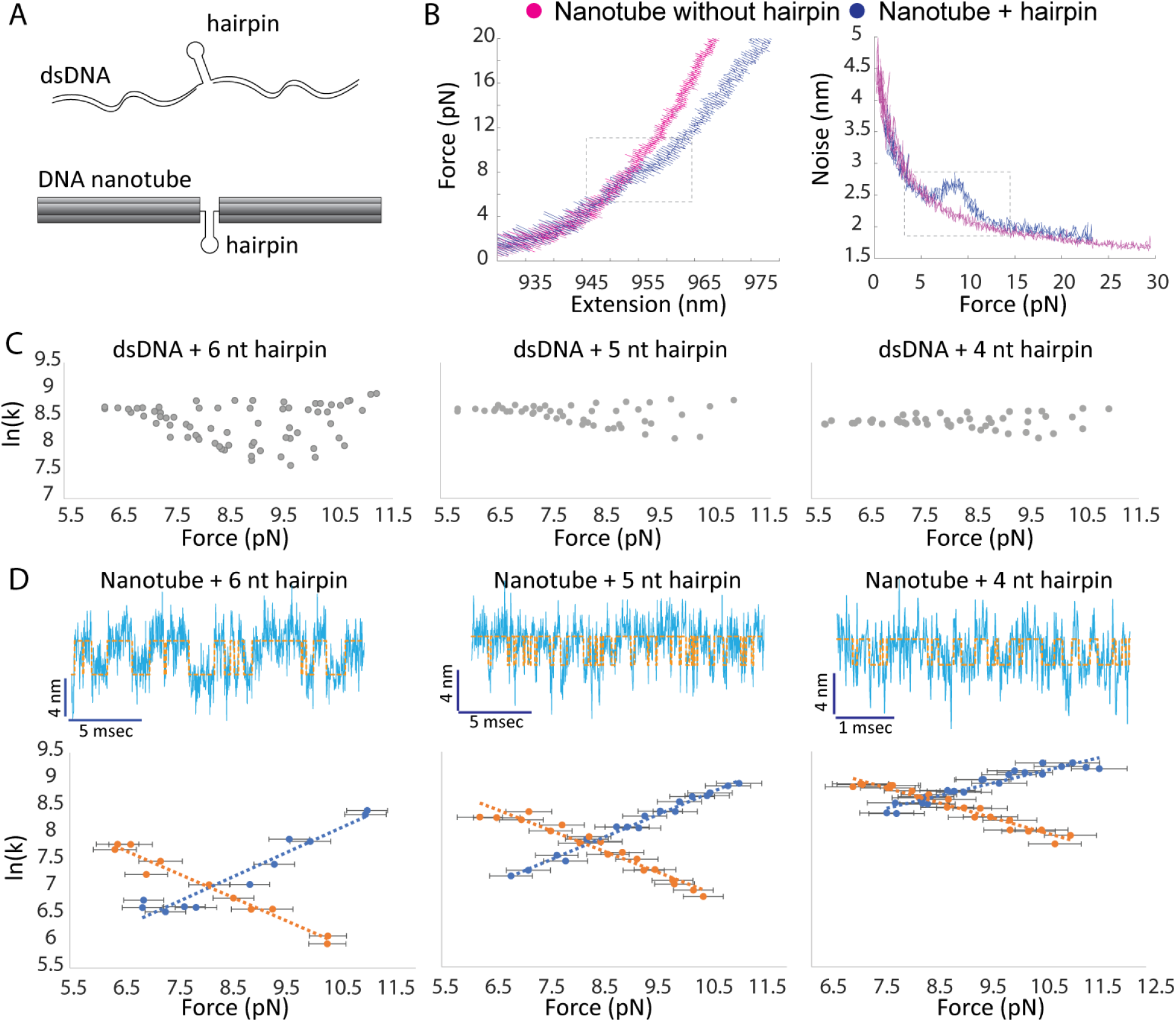
Direct visualization of the folding and unfolding kinetics of short DNA hairpins using DNA nanotube tethers. A) 6, 5, or 4 nt DNA hairpins are introduced in nanotube or dsDNA tethers. The other end of the nanotubes are attached to the optically trapped beads. B) left panel: force extension curves of DNA nanotubes with and without DNA hairpin. The extension curve with hairpin shows a clear shoulder in the range of 6 - 10 pN. Right panel: amplitude in extension noise plotted against force from the data shown in the left panel. C) Analysis of hairpin folding and unfolding using dsDNA tethers. The rates were extracted from dwell time analysis of states found by HMM. D) Characterization of hairpin folding and unfolding using nanotube tethers. Top panels: raw extension data is shown in blue, states assigned by HMM are shown as dotted orange lines, the force applied in these plots are about 8 pN. Bottom panels: force dependent folding and unfolding rates extracted from dwell time analysis. n=3 for all three DNA hairpins shown. Error bars represent standard deviation.

## Conclusion

In this study, we systematically evaluated how noise is distributed across different frequencies in dual trap optical tweezers systems, and showed that rigid DNA nanotube tethers only suppress high frequency noise when compared to dsDNA. This suggests that the use of rigid tethers is advantageous when studying single molecule events that have dwell times in the sub-millisecond time scale, while there are no obvious advantages in using the rigid tethers when studying slower events. Indeed, Pfitzner et al.3 demonstrated that when extracting folding kinetics of a slow-folding, long DNA hairpin, both dsDNA and rigid DNA nanotube tethers gave near identical kinetics results, while according to both our data and the data report by Pfitzner et al., the folding and unfolding of short DNA hairpins can only be directly observed when using rigid nanotube tethers. Using our rigid nanotube system, we showed that we can directly observe and resolve the fast-folding kinetics of a 4 nt DNA hairpin, which has a transition of about 2.5 nm and average dwell time in the hundreds of microseconds time scale. This is the smallest and fastest transition event that has been directly observed via optical tweezers. We believe that rigid DNA nanotube linkers can be a powerful tool to study fast folding events of small biomolecules^18^ or capturing folding transition paths^19,20^, which can occur in the microseconds time scale.

## Appendix A Derivation of GLE

To prove Eq. 4, we first switch into an alternate coordinate system, *x*(*t*) = *x*_1_(*t*) – *x*_2_(*t*) and *y*(*t*) = *x*_1_(*t*) + *x*_2_(*t*). Replacing these in Eq. 1, we get:

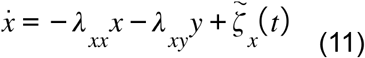

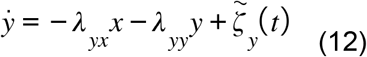

where

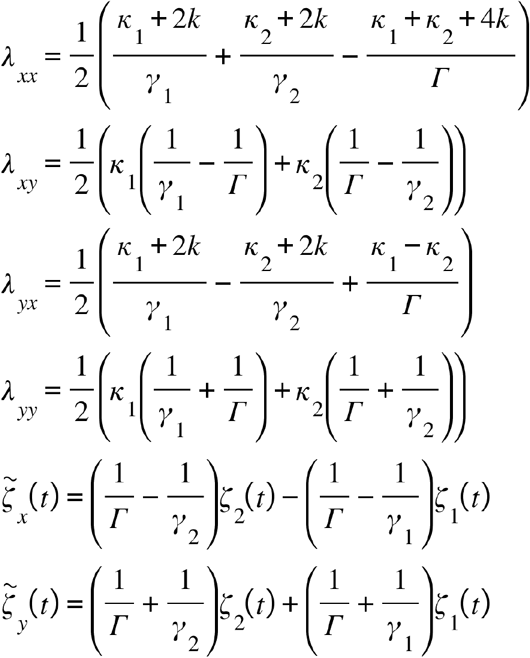

Eq. 12 is a linear ordinary differential equation with the following general solution:

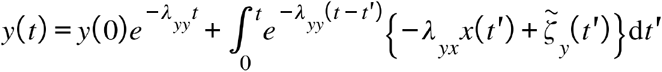

Evaluating the integral containing *x*(*t*) by parts, we get:

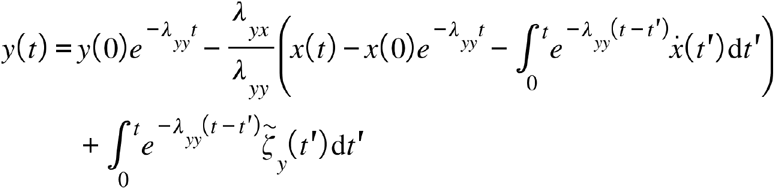

Plugging this equation for *y*(*t*) into Eq. 11 and taking the limit 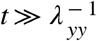, we arrive at the following GLE for *x*:

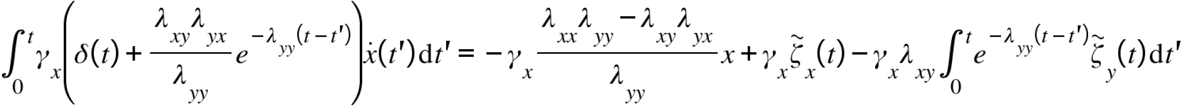

This is equivalent to Eq. 4, such that:

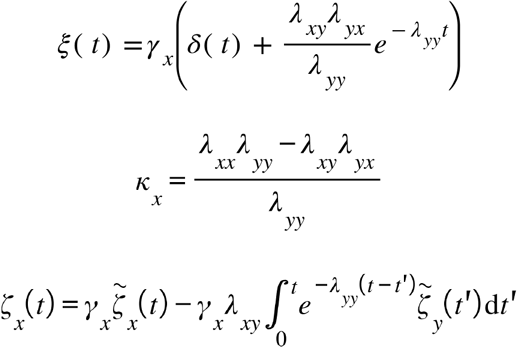

and 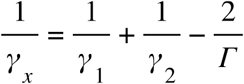 is a characteristic friction coefficient along *x*. To prove Eq. 5, we first show that the first moment of the stochastic force is zero:

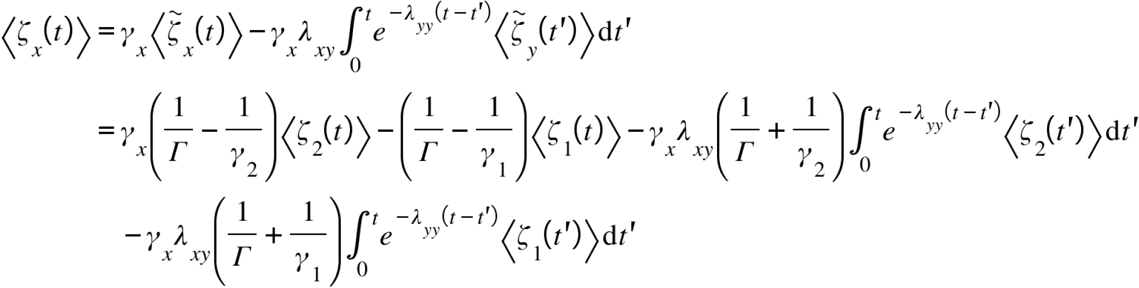

where we used Eq. 3 to compute the averages in the last statement. Similarly, the second moment of the stochastic force along *x* can be shown to be related to the memory kernel:

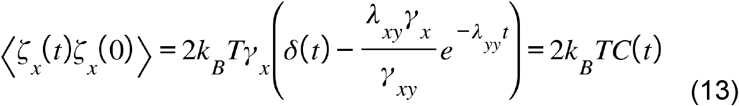

where 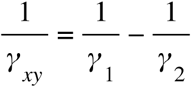 is another characteristic friction of the system. Note that both *ξ*(*t*) and *C*(*t*) have a general form, as in Eq. 6, given by a sum of two terms: a fast decaying Markovian component and an exponential term with a finite decay time 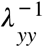 and a coefficient *λ′*. Thus, the bead to bead extension is generally a non-Markovian coordinate with finite memory that depends on the parameters of the traps, the beads, and the tether: *κ*_1_, *κ*_2_ *k*, *γ*_1_, *γ*_2_, and *Γ*.

## Appendix B Derivation of PSD

To calculate the PSD for a time series *x*(*t*) that follows the GLE (Eq. 4), we first take the Fourier Transform on both sides:

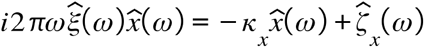

Rearranging the terms, we get:

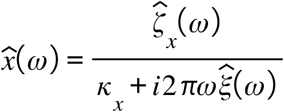

Taking the expectation of the square of the absolute value on both sides and dividing by the total measurement time *T*, we get:

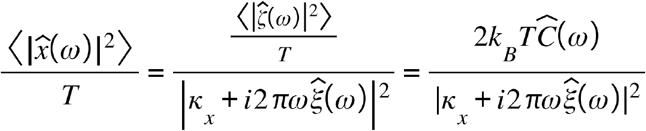

where we used Eq. 13 in the last statement.

## Materials and Methods

### Synthesis of DNA origami nanotubes

The 6 helix bundle DNA origami nanotube was designed with caDNAno software [6]. ssDNA scaffold (p8064) was purchased from tilibit nanosystems (Munich, Germany), and oligonucleotide staples along with biotin and digoxigenin modified oligos were purchased from integrated DNA technologies (San Diego, USA). The 6 helix bundle was assembled using one pot synthesis: 20 nM of p8064 was mixed with 140 nM staples (140 nM for each staple) in 1X folding buffer (5 mM Tris (pH 8.5), 1 mM EDTA, 12 mM MgCl2). The mixture was heated up to 65oC for 10 mins, followed by slow cooling to 25oC over 14 hrs. Excess staples were removed by repetitive concentration-dilution in Amicon 100k MWCO (Merck Millipore, USA) in 1X folding buffer. The digoxigenin and biotin modified nanotubes were mixed at a 1:1 ratio at a final concentration of 10 nM in 1X folding buffer and placed at rt overnight to allow dimerization.

### Preparation of dsDNA tethers

dsDNA tether was prepared by PCR on lambda phage DNA using the following primers: gtggaatgc-catgtgggctgtcaa and gattaccgtaagacggaaatcactccc.

### Characterization of folded DNA nanotubes

#### Agarose gel electrophoresis

2% agarose gel was prepared in 1X TBE buffer supplemented with 10 mM MgCl2 and prestained with Sybr safe (Thermofisher, USA). 5 nM 10 ul DNA nanotube was mixed with DNA loading dye (Thermofisher, USA). The gel was run at 90 V in the cold room with prechilled 1X TBE supplemented with 10 mM MgCl2 for 45 mins.

#### AFM imaging of the 6 helix bundle

The folded and purified 6 helix bundle was adjusted to 5 nM in 1X folding buffer. 10 ul of the nanotube solution was pipetted onto freshly cleaved mica, and after 5 mins NiSO4 was added to a final concentration of 2 mM and incubated an additional 10 mins. The surface was washed with 1 ml of 1X folding buffer and imaging was carried out in 1X folding buffer using the Bruker Multimode 8 with Peakforce-HIRS-F-B tips and tapping mode. AFM data was processed using a combination of Gwyddion and ImageJ.

#### Optical tweezers measurements

The instrument used in this study is a home built dual trap optical tweezer system, which is described in detail in [12]. Briefly, an Nd:YAG 1,064 nm laser was used and an acousto optic deflector (AOD) was used to switch the single laser beam every 5 μsec, thus generating two traps. Detection of bead position was measured by monitoring scattering of laser on a quadrant photodiode (QPD). Streptavidin modified beads were directly ordered from bandlabs (catalog number: CP01004) and passivated by incubation in BSA solution (1 mg/ml in TE 40 buffer: 40 mM Tris pH 8.0, 1 mM EDTA) for 1 hr with constant shaking and washed by repetitive centrifugation (5 min 4500 rcf) and resuspension in TE40 buffer. Finally the beads were resuspended in TE40 at a concentration of 0.2% (w/v). Anti-digoxigenin antibodies were coupled to 1 um polystyrene beads via NHS-EDC coupling: 50 ul of 5% (w/v) 1 um carboxylated polystyrene beads (Spherotech, USA) was buffer exchanged in to reaction buffer (MES-NaOH, pH6, 500 mM NaCl) via repetitive centrifugation (3000 rcf 3 mins) - resuspension, the final concentration of the beads was adjusted to 5% (w/v). EDC (1-ethyl-3-(−3-dimethylaminopropyl) carbodiimide hydrochloride) and Sulfo-NHS (Sulfo-NHS, Hydroxy-2,5-dioxopyrrolidine-3-sulfonicacid sodium salt) was added to the bead solution with a final concentration of 5 mM and 10 mM. The reaction was incubated at room temperature for 15 mins and 2.8 ul of 40 mM beta mercaptoethanol was added to quench the reaction. Afterwards, Anti-digoxigenin antibodies were added to the beads and incubated at room temperature for 2 hr and constant shaking. The antibody-modified beads were then washed with TE40 via repetitive centrifugation (3 min 3000 rcf) - resuspension and adjusted to a final concentration of 0.2Loading the DNA nanotube tether onto anti-digoxigenin beads: nanotubes from the dimerization reaction was adjusted to 0.5 nM, and 3.2 ul of it was added to 3.2 ul of 0.2% anti-DIG beads. After incubation at room temperature for 15 mins, the beads were diluted to 900 ul with TE40.

## Supplementary Information

**Figure S1:**
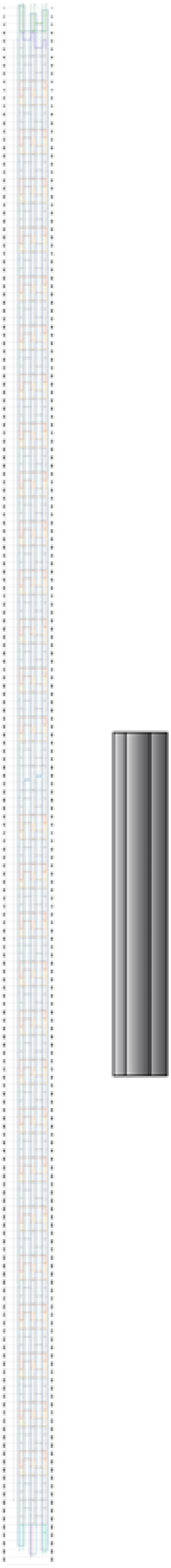
Left panel: caDNAno design scheme of the 6 helix bundle. Green oligos are either modified with digoxigenin or biotin. Right panel: 3D rendering of the 6 helix bundle using Autodesk Maya.

**Figure S2:**
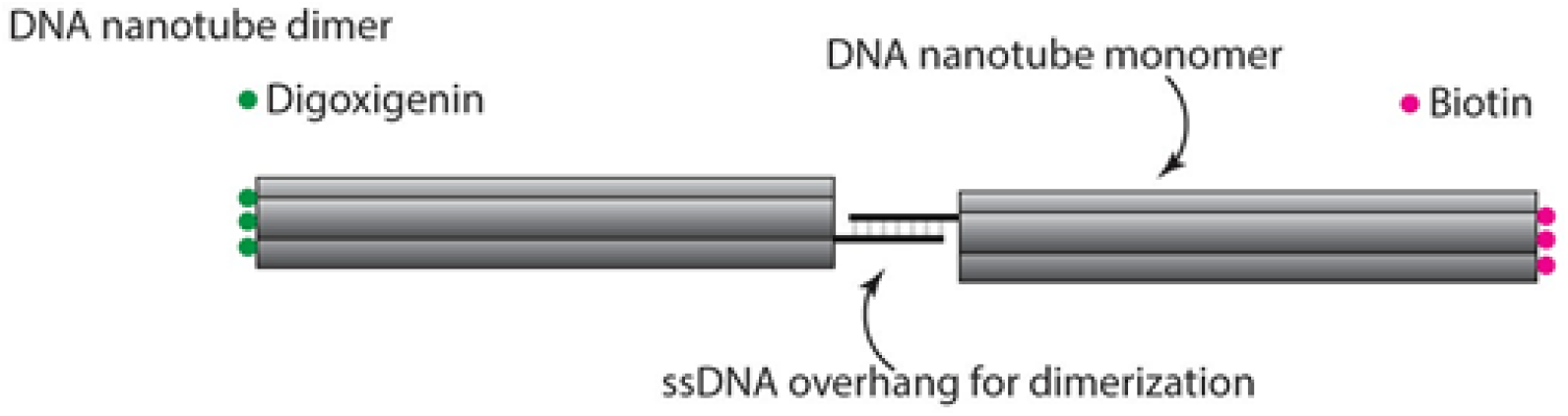
Design scheme for generating DNA nanotube dimers. Digoxigenin and biotin modified nanotube monomers are dimerized via DNA hybridization.

**Figure S3:**
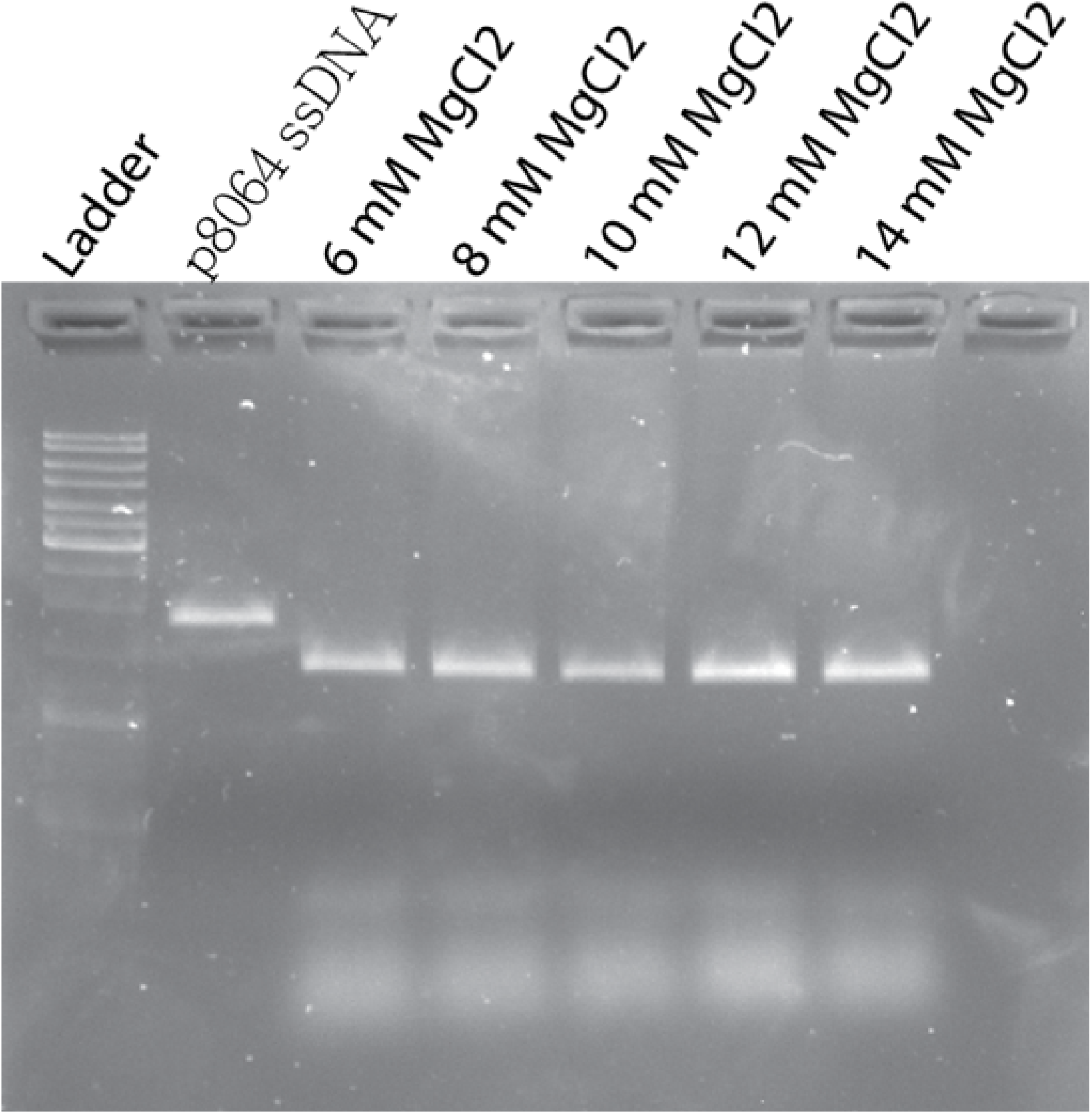
2% agarose gel pre-stained with Sybr Safe showing successful folding of the 6 helix bundle at various magnesium concentrations. 12 mM MgCl2 was chosen for optical tweezers experiments and AFM imaging.

**Figure S4:**
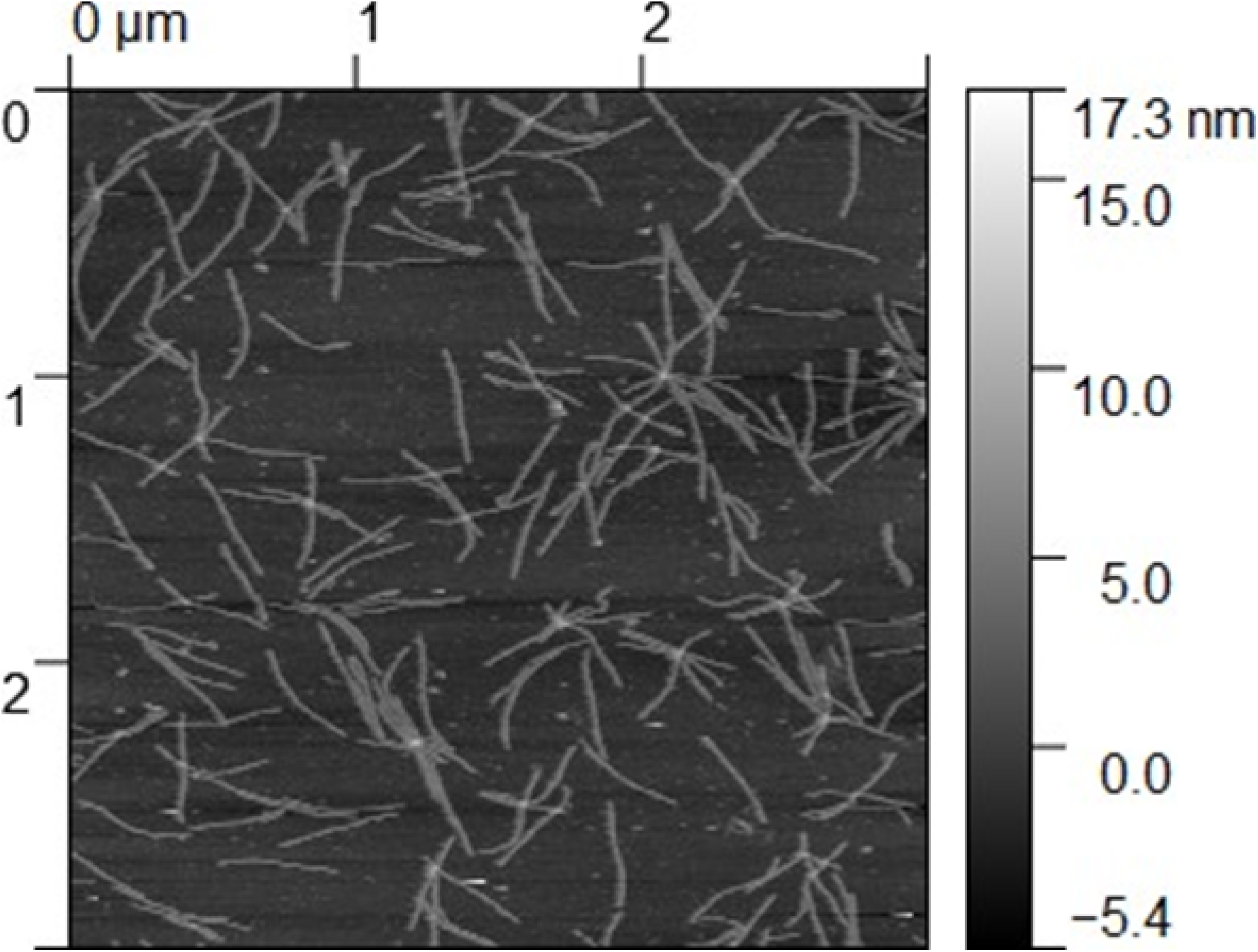
AFM micrograph of purified 6 helix bundle DNA nanotube imaged with a 2000 nm x 2000 nm window. Raw data was processed with Gwyddion.

**Figure S5:**
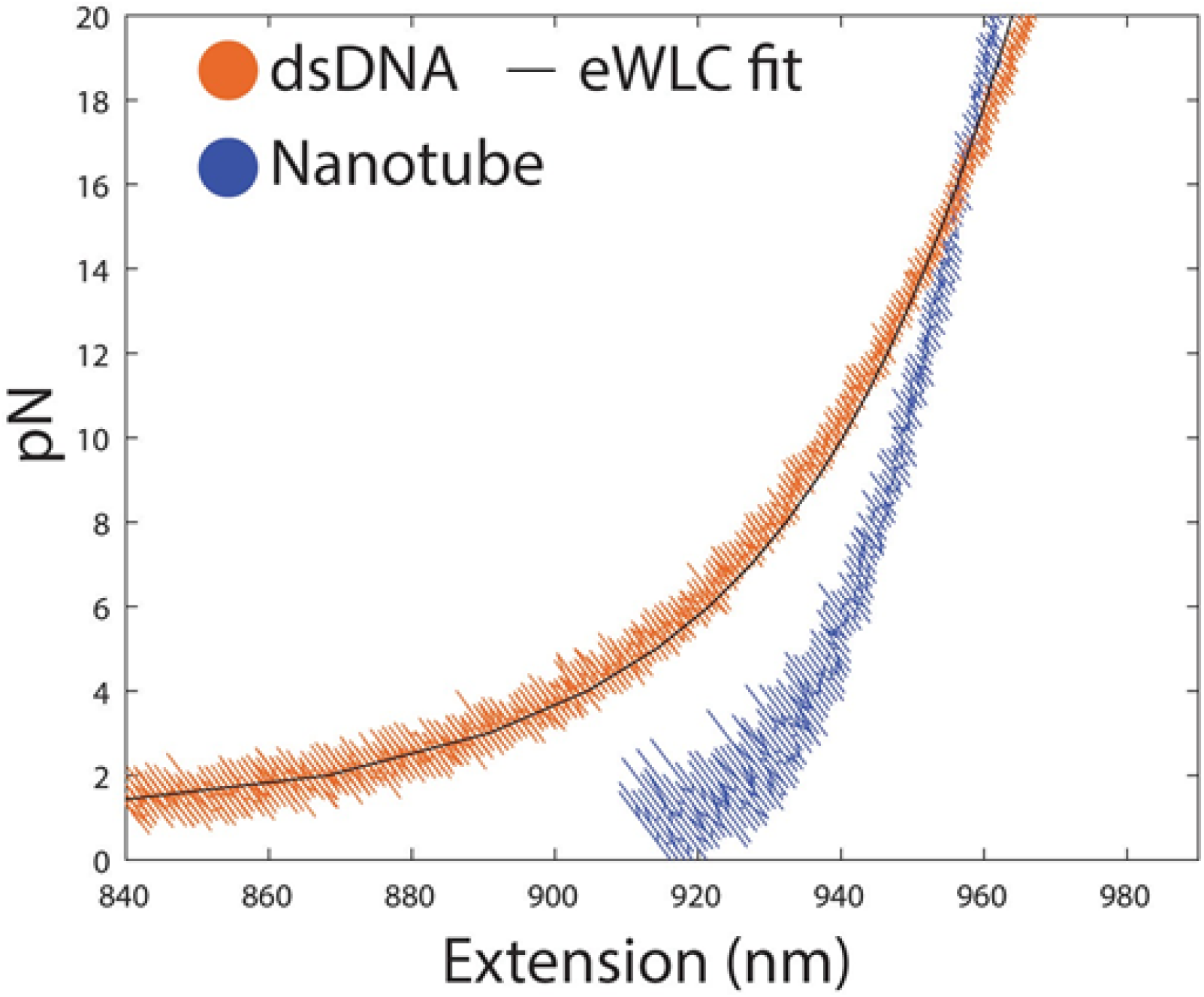
Force extension curves of dsDNA tether and nanotube tether. The force extension curve of dsDNA fits well with the extensible worm-like chain model (persistence length = 35.6 nm, contour length = 985.0 nm).

**Figure S6:**
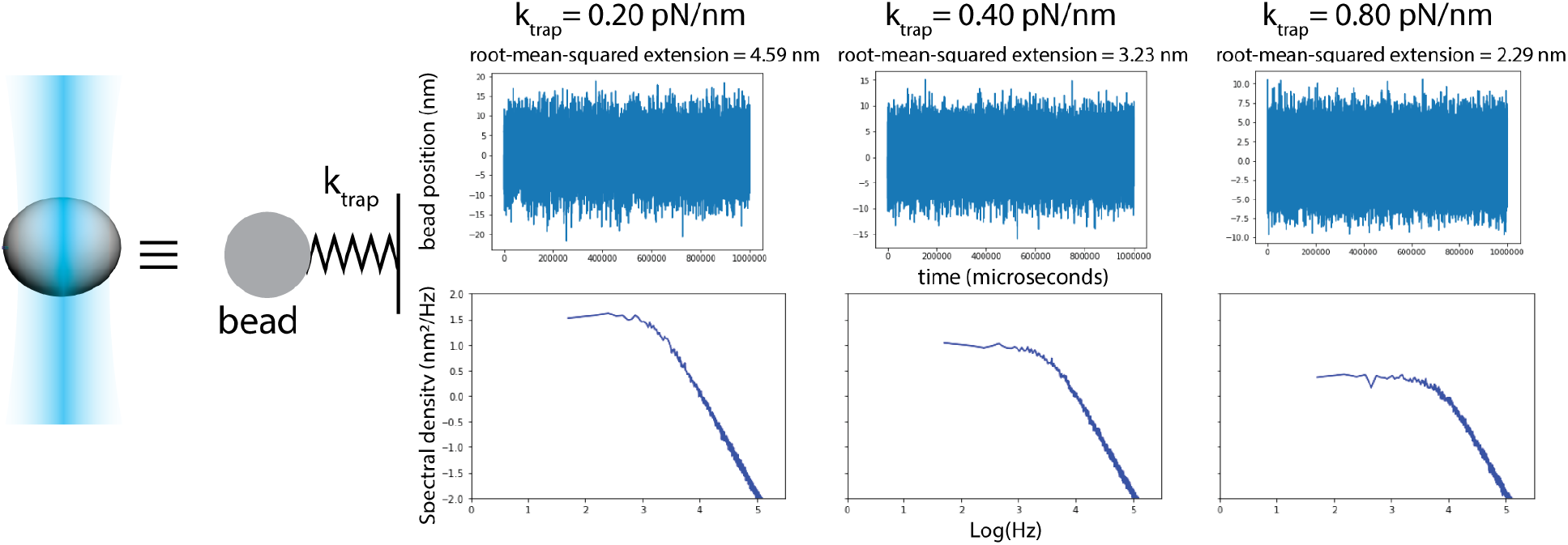
Simulating one optically trapped bead with different trap stiffness. The raw simulated data for bead position over time is plotted in the top panel. The power spectrum (bottom panel) exhibits Langevin dynamics. The root-mean-squared extension (noise) decreases as the trap stiffness increases. The area under the PSD represents noise, and from the plots we can clearly observe that as we start to downsample the data (integrating the area under the PSD up to a certain frequency), the noise from the soft trap will always be higher than that of stiffer traps.

